# Changes in Corticospinal Excitability in Response to Mediolateral Gait Instability

**DOI:** 10.1101/2025.01.30.635462

**Authors:** Raven O. Huiberts, Sjoerd M. Bruijn, Jennifer L. Davies

## Abstract

Unpredictable gait disturbances, particularly in the mediolateral direction, pose a significant challenge to stability and are a common contributor to falls. Although the corticospinal tract is critical for gait and postural control, its response to such instabilities remains unclear. To investigate if corticospinal excitability increases during laterally destabilised gait, single-pulse transcranial magnetic stimulations were delivered over the primary motor cortex of 15 healthy individuals during steady-state and laterally destabilised treadmill gait. Full-body kinematics were recorded using an optoelectronic motion capture system. Stimulations with coil displacement >5 mm from the targeted location were excluded. Corticospinal excitability was quantified for four upper- and three lower-leg muscles by the motor evoked potential (MEP) amplitude and compared between steady-state and destabilised gait. Destabilisation resulted in a wider step width and shorter stride duration with increased variability and greater dynamic instability. Foot placement control was increased at mid-swing, along with greater average foot placement error. No differences in corticospinal excitability were observed in the lower-leg muscles. All upper-leg muscles demonstrated greater absolute MEPs in destabilised relative to steady-state gait. After normalising MEP to the pre-stimulus muscle activity, these periods became less pronounced, however, increases were observed in all but the gastrocnemius muscles. These findings suggest heightened readiness of the corticospinal tract projecting to upper-leg muscles during destabilised gait, which could reflect general stabilising strategies such as decreasing stride time and increasing step width.

**KEY POINTS:**

- Maintaining stability during bipedal gait requires precise control, especially when confronted with unpredictable mediolateral disturbances, a common contributor to falls.
- When walking on a treadmill that was laterally destablised, participants adopted a cautious walking strategy characterised by wider step width, shorter stride duration and increased variability.
- Corticospinal excitability to upper-leg muscles was greater in destabilised than steady-state gait, corticospinal excitability to lower-leg muscles remained largely unchanged.
- The observed increases in corticospinal excitability to upper-leg muscles likely reflect a general stabilisation strategy in response to a laterally destabilised walking surface.
- Increases in MEP gain (MEP amplitude relative to ongoing muscle activity) suggest that changes in corticospinal excitability may reflect additional roles beyond modulating the ongoing muscle activity.

## INTRODUCTION

Human gait is a complex task that requires coordinated activity of around 200 muscles. Despite the complexity of human gait, it can be seemingly effortless. Historically, it was assumed that this was achieved primarily through spinal control systems, a notion based on observations that quadrupedal mammals are capable of basic locomotion rhythmicity even in the absence of supraspinal processing [1–6]. While there are shared control mechanisms between human and quadrupedal gait [7,8], cortical lesions in humans can drastically debilitate gait [9,10] indicating an important role for supraspinal control. This likely reflects the greater stability demands of bipedal gait [11]. Understanding these supraspinal processes is essential, particularly for advancing fall prevention strategies and neurorehabilitation.

Transcranial magnetic stimulation (TMS) can be used to probe the corticospinal pathway. TMS over the motor cortex depolarises pyramidal tract cells and, with sufficient excitation, this results in the generation of action potentials that descend along the corticospinal tract to depolarise alpha motor neurones in the spinal cord [12–14]. The subsequent firing of motor units can be measured using surface electromyography (EMG) as a motor evoked potential (MEP). The size of the MEP reflects the responsiveness of motor cortical circuitry, segmental motor circuits and alpha motor neurones [15–17], which we will refer to as corticospinal excitability.

Using TMS, it has been demonstrated that corticospinal excitability changes throughout the gait cycle, generally aligning with phasic muscle activity [17–20]. However, these excitability patterns do not always mirror ongoing muscle activity. Schubert and colleagues [21] observed increased corticospinal excitability prior to phasic muscle activation and suggested that this reflects increased supraspinal input as an anticipatory adjustment aimed to preserve gait stability.

Gait stability is the ability to counteract the forces that induce moments to accelerate our centre of mass (CoM) away from the base of support, such as gravity and internal and external perturbations, by generating appropriate muscles force in order to remain upright [22–24]. This requires active control, particularly in the mediolateral direction, where passive dynamics offer limited support [25,26]. To maintain mediolateral stability, the body employs several strategies to either redirect the CoM through stance leg control or adjusting the base of support at foot placement to accommodate CoM trajectory, which happens through the swing-leg mechanics [11,27–30]. These methods of maintaining mediolateral stability involve both ankle crossing muscles [11,31,32], which contribute to redirecting the CoM, and hip crossing muscles, primarily the gluteus medius, which are involved in controlling foot placement [27,28,33–35].

Mediolateral gait stability deteriorates with ageing [36,37], which manifests as reduced local dynamic stability and increased gait variability [38–40]. These changes are well-established indicators of an increased risk of falling [41–45]. Improving our understanding of how mediolateral gait stability is maintained is essential in the development of strategies to prevent and reverse this deterioration, thereby reducing risk of falls.

Human gait studies have shown increased corticospinal excitability with precise stepping [46,47] and external mechanical gait constraints [48]. These findings suggest corticospinal contribution increases with greater gait control demands. Camus et al. [49] demonstrated that humans can adaptively modulate corticospinal excitability in anticipation of unpredictable central perturbations during walking, supporting the idea proposed by Schubert et al. [21] that supraspinal input can serve an anticipatory adjustment to preserve gait stability. Furthermore, during standing balance, corticospinal excitability of the tibialis anterior muscle scales with mediolateral sway velocity, further implying a role of the corticospinal tract in stability control [50]. However, it is not known if there is modulation of corticospinal excitability of leg muscles during mediolaterally unstable gait, particularly when the instability is unpredictable (both in timing and direction), such as missteps or lateral pushes in daily life. This is important to understand, as falls often result from unpredictable mediolateral disturbances.

Here, we test if corticospinal excitability increases when walking on a mediolaterally destabilised treadmill. We hypothesise that corticospinal excitability will increase in accordance with the mediolateral stability demands. We expect that increases in corticospinal excitability will be observed throughout the gait cycle, reflecting the heightened readiness to respond to unpredictable perturbations.

## METHODS

### Participants

Twenty-three participants (seven male, 16 female; aged 18–40 years) provided written informed consent to participate. All participants reported that they were healthy adults without any contraindications to TMS or other risk factors that could impact the safety or reliability of this study (https://doi.org/10.17605/OSF.IO/7AJUZ and [51]). Participants were informed that they could withdraw from the experiment at any time. All procedures were conducted in accordance with the Cardiff University School of Healthcare Sciences Research Ethics Committee. There were no adverse events. Six participants were excluded because no MEP could be observed within a comfortable stimulation intensity. One participant was excluded because they had to leave early. After processing, another participant was excluded because the TMS coil moved too much during the walking trials, resulting in less than 100 useful stimulations per condition. Data from the remaining 15 participants are reported here.

### Procedures

The participants performed both steady-state and laterally destabilised treadmill gait within a Gait Realtime Analysis Interactive Lab (GRAIL) system (Motek Medical BV, Houten, Netherlands). Destabilisation was induced by continuous pseudorandom mediolateral oscillations of the treadmill with a peak-to-peak amplitude of 100 mm (Figure 1 C). The velocity of the treadmill was set at a constant speed of 4.0 km/h. Each condition (steady-state and laterally destabilised) consisted of a 200-s familiarisation period followed by a 20-min trial. The two conditions were performed in a single session with a break in between. The order of the conditions was randomised.

**Figure 1.**
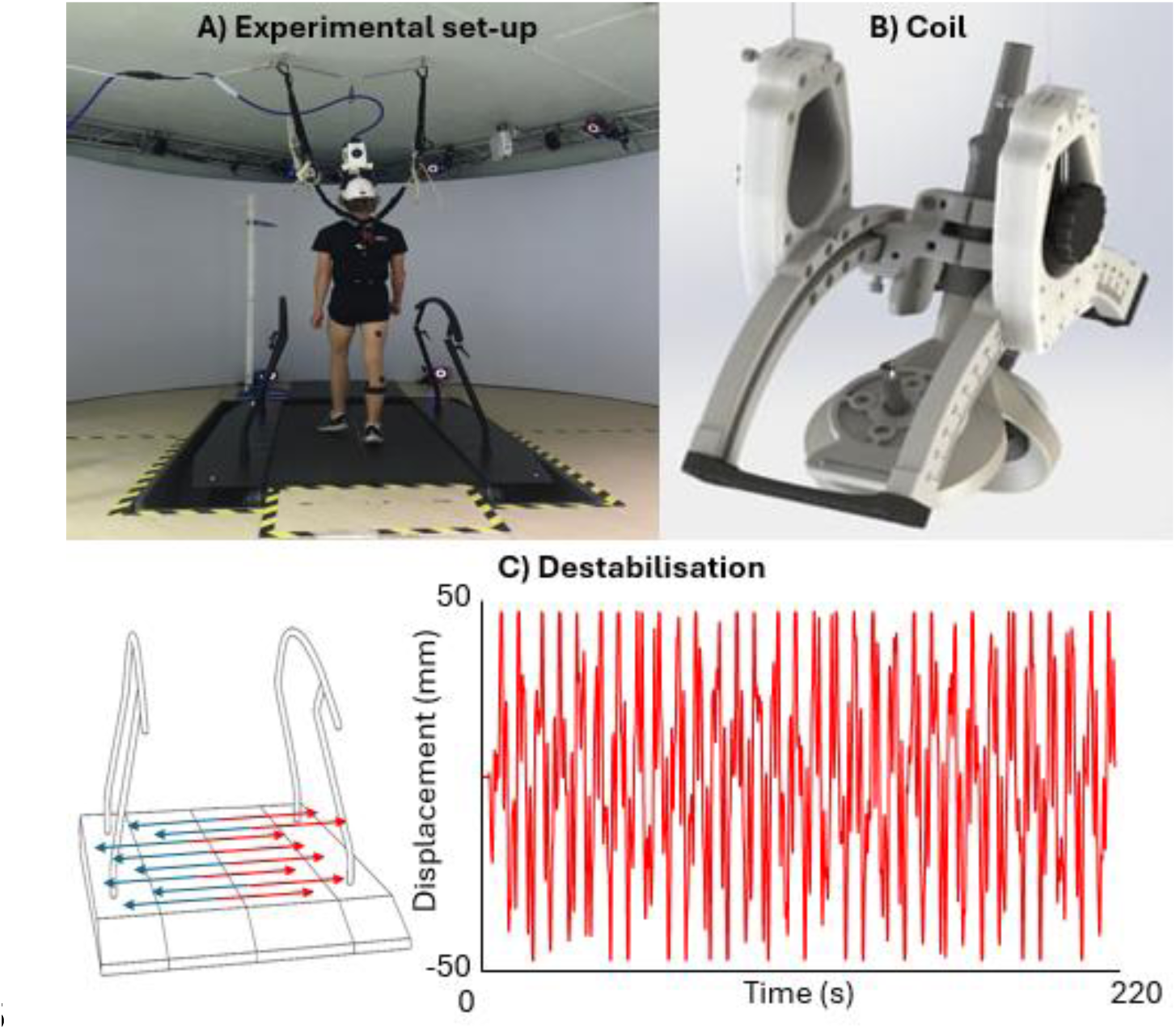
Display of the experimental set-up. A) Overview of the full experimental setup. B) Close-up view of the batwing coil used for stimulation. C) Illustration of the lateral destabilisation mechanism, implemented via treadmill surface shifts. The accompanying graph shows the time course of lateral displacement

### Motion capture

During the familiarisation and the trials, full-body kinematics were captured at 100 Hz using Vicon Nexus software (version 2.12, Oxford, UK). Spherical retroreflective markers (12-mm diameter) were placed in accordance with the full-body Plug-in Gait model (Vicon 2.12, Oxford, UK) with some modifications: the markers on the side/back of the head were placed on the side/back of the helmet (Figure 1 A & B), and additional markers were placed on the nasal bridge and the left and right tragus.

For real-time detection of heel strike, the HBM 2.0 model (Motek Medical BV, https://knowledge.motekmedical.com/wp-content/uploads/2019/07/HBM2-Reference-Manual-Full-Body.pdf) was used. Kinematic data were streamed into D-flow software (version 3.36, Motek Medical BV) and processed in real time.

### EMG

Before the trials, surface electrodes (Trigno Avanti Sensor, Delsys, Boston, MA, USA) were placed on the following muscles on the right leg according to the SENIAM guidelines ([SENIAM](http://www.seniam.org/)): adductor magnus, biceps femoris, gluteus medius, rectus femoris, tibialis anterior, gastrocnemius medialis, gastrocnemius lateralis. EMG data was collected at a sampling frequency of 2000 Hz in Vicon Nexus software and was synchronised with kinematic data.

### TMS

TMS (posterior-anterior current) was administered using a custom-made bent batwing coil (Magstim, Whitland, Wales). The coil was positioned on the scalp over the left motor cortex and position was adjusted until MEPs could be observed in the tibialis anterior of the right leg. The tibialis anterior was used for this purpose because it exhibits a clear, easily measurable response [20], allowing us to confirm when the stimulation was being delivered to the leg area of the primary motor cortex. Our previous work indicates that individual leg muscles have overlapping cortical representations, such that several muscles can be stimulated from one location on the scalp [52]. It also indicates that individual leg muscles have multiple ‘hot spots’, such that there is not one single optimal location for each muscle [52]. Although the position was not optimised for each individual muscle, we were able to detect MEPs from multiple muscles during gait from stimulation at this single location on the scalp.

The position of the coil on the head of the participant was fixed using of a custom-designed helmet (Figure 1). The weight of the coil and helmet was supported by two spring-coil balancers that were attached the ceiling over the centre of the treadmill. The tension in the spring-coil balancers was adjusted for each participant so that the coil rested on the scalp with enough weight to ensure stability of coil position throughout the gait cycle (i.e., to prevent the coil ‘bouncing’ with vertical head movement) but enough weight removed so that this would not become uncomfortable over the duration of the experiment.

For each participant, motor threshold was determined as the minimal stimulation intensity required to evoke an MEP in tibialis anterior while the participant was standing quietly. The threshold was determined according to the following procedure: First, the intensity of the stimulator was set to 35% of the maximal stimulation output, which (for most individuals) is a subthreshold intensity within the limits of tolerability. Thereafter, the intensity was progressively increased in steps of 5% until a discernible MEP could be observed in at least 5 out of 10 stimulations, or the participant reported that they found the stimulation intolerable. If the latter occurred, the session was terminated (n = 6/23 participants). If the former occurred, the intensity was decreased in steps of 2% until the simulation did not evoke an MEP in 5 out of 10 stimulations. Lastly, the intensity was increased again in steps of 1% until the lowest intensity that consistently evoked a MEP was found.

During the experimental trials, the TMS intensity was set at 110% of this motor threshold. Stimulations were given with random delays after heel strike with a varying number of strides between the stimulations (ranging from 3 to 5 strides). This resulted in approximately 200 stimulations over the 20-minute trial, evenly distributed across the stride cycle. The stimulation instant was recorded from the output of Dflow via a Phidget Interface Kit 8/8/8 (Phidgets Inc. Calgary, AB, Canada) with a Trigno Analog Input Adaptor (Delsys) that was sampled at 2000 Hz in Vicon Nexus software (version 2.12) synchronised with EMG data.

### Relative coil displacement

Prior to the experiment, a static measurement was conducted using an additional marker placed at a central point within the helmet. During the experimental trials, the virtual marker’s position was calculated using both the fixed helmet markers and the head markers. To determine the displacement of the helmet relative to the head, the deviation of the virtual marker calculated from the head markers was compared to its intended position, as defined by the rigid markers on the helmet.

### Data processing

Gait events for offline analysis were calculated with a custom code where heel strike was identified as the moment the heel marker reach its lowest point and toe-off was identified as the moment the heel marker reached its highest upwards acceleration (https://github.com/SjoerdBruijn/GaitEvents).

#### Gait pattern

The gait pattern in each condition was described using step width (average and variance), stride duration (average and variance), dynamic stability and quality of foot placement control in the last 150 strides of the 200-s familiarisation period.

Step width was defined as the mediolateral distance between the heels at heel strike. Stride duration was defined as the time interval between two consecutive right heel strikes.

Dynamic stability was calculated with the local divergence exponent (λ_s_), a measure that quantifies the resilience to infinitesimal perturbations that occur during gait [53]. It is computed by determining the average exponential rate of divergence of neighbouring trajectories in state space [54]. Higher values indicate greater local instability. For these calculations, Rosenstein’s algorithm was used [55]. The state space was created from time-normalised mediolateral velocity time series of the averaged thorax markers (C7 and left and right acromion) and five time-delayed copies. The λ_s_ was estimated as the slope of the mean divergence curve, with a normalised stride time from 0 to 0.5 stride [56,57].

The quality of foot placement control was calculated using the model of Wang and Srinivasan [28] (utilising codes from [58]). This model assumes a linear relation between the deviations in CoM state and subsequent foot placement (Equation below). Within our analysis, the pelvis (PEL), defined as the mean of the right and left posterior superior iliac spine markers, was used as a proxy of the CoM [59]. The foot placement states were de-meaned to allow for interpretation.

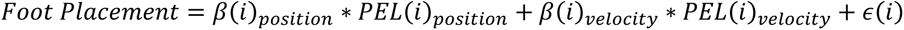

- *β_positin_* = *position regression coefficient*
- *β_velocity_* = *velocity regression coefficient*
- *ϵ* = *residual variance* (i.e. foot placement error)

The foot placement was calculated in the single-leg stance phase, defined as the time between contralateral toe off and heel strike. To account for displacement, such as wandering backward, forward, left, or right, the coordinate system was taken from the previous stance foot. Additionally, within the laterally oscillating treadmill condition, the PEL_velocity_ was corrected by adding the treadmill’s lateral velocity to the derivative of the PEL_position_. The degree of foot placement control was quantified by the relative explained variance (R^2^). To attain normality in the data fisher transformation was performed on the R^2^ values. Lastly, as the R^2^ indicates the percentage of the foot placement variance explained by the variance in PEL states, we also calculated the average foot placement error as a measure of foot placement precision. This was calculated as the standard deviation of ϵ at the instant of foot placement.

#### Ongoing EMG and MEPs

EMG signals were bandpass filtered between 20 Hz and 500 Hz using a second-order bi-directional Butterworth filter. Ongoing EMG was calculated by rectifying and then averaging the filtered signal from 20 ms prior to stimulation to 10 ms after stimulation. The stimulation artifact was excluded by removing the data point at the time of stimulation (Figure 2).

**Figure 2.**
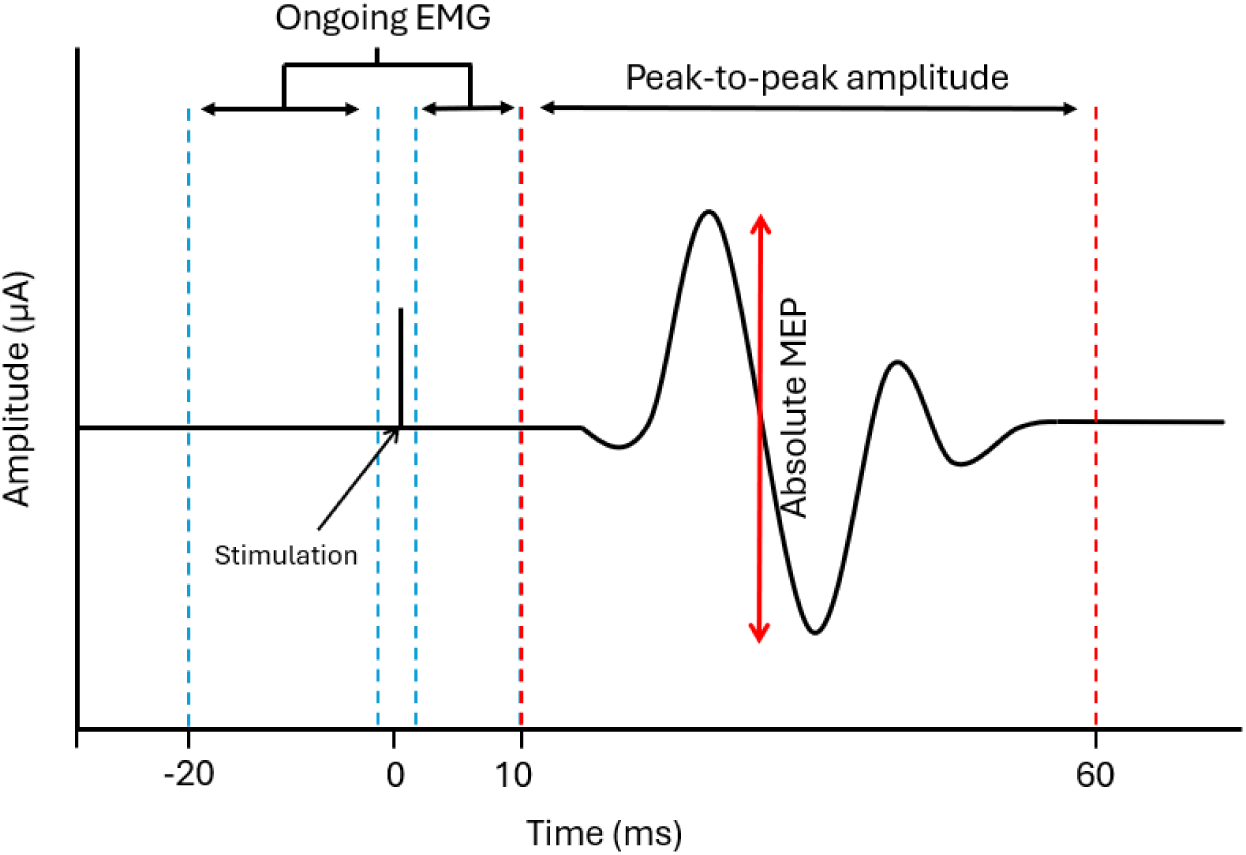
Calculation of ongoing EMG and absolute MEPs. The blue dotted lines indicate the windows over which the data was rectified and averaged to calculate the ongoing EMG. The red dotted lines indicate the window over which the peak-to-peak value was calculated indicating the absolute MEP. MEP gain was calculated by dividing absolute MEP amplitude by ongoing EMG.

Absolute MEP amplitudes were calculated as the peak-to-peak amplitude of the filtered signal within the time window of 10 to 60 ms post-stimulation. MEP gain was calculated by dividing the absolute MEP amplitude by the ongoing EMG (Figure 2). MEPs were excluded if the relative helmet-head position at the time of stimulation was >5 mm along the vertical axis or >10 mm in the horizontal plane (combined displacement) from the original relative position. For each stimulation, the corresponding percentage of the gait cycle was determined based on the percentage of the total time between consecutive right heel strikes.

For the ongoing EMG, absolute MEPs and MEP gains, all data points were smoothed, utilising SMART [60], with adjustments. First, all data points were linked to their corresponding gait cycle percentage (Figure 3A). Thereafter, the estimated time-series was reconstructed for each participant by convolving the kernel with the outcomes. This was then divided by the sum of the kernel density (weights) to maintain the proper scaling (Figure 3B & C). After this, two separate methods were performed: a weight-based approach and a non-weight-based approach. For the weight-based approach, normalised weights were calculated and averaged across participants for each time point in the gait cycle. This method ensures that participants with higher sample density at specific percentages of the gait cycle have a greater impact on the smoothed outcome at those corresponding time points. However, while this takes into account data distribution, it can also cause slight misinterpretation due to overrepresentation of individuals at different timepoints. Therefore, we also calculated the average across participants for each time point without any weight adjustment.

**Figure 3.**
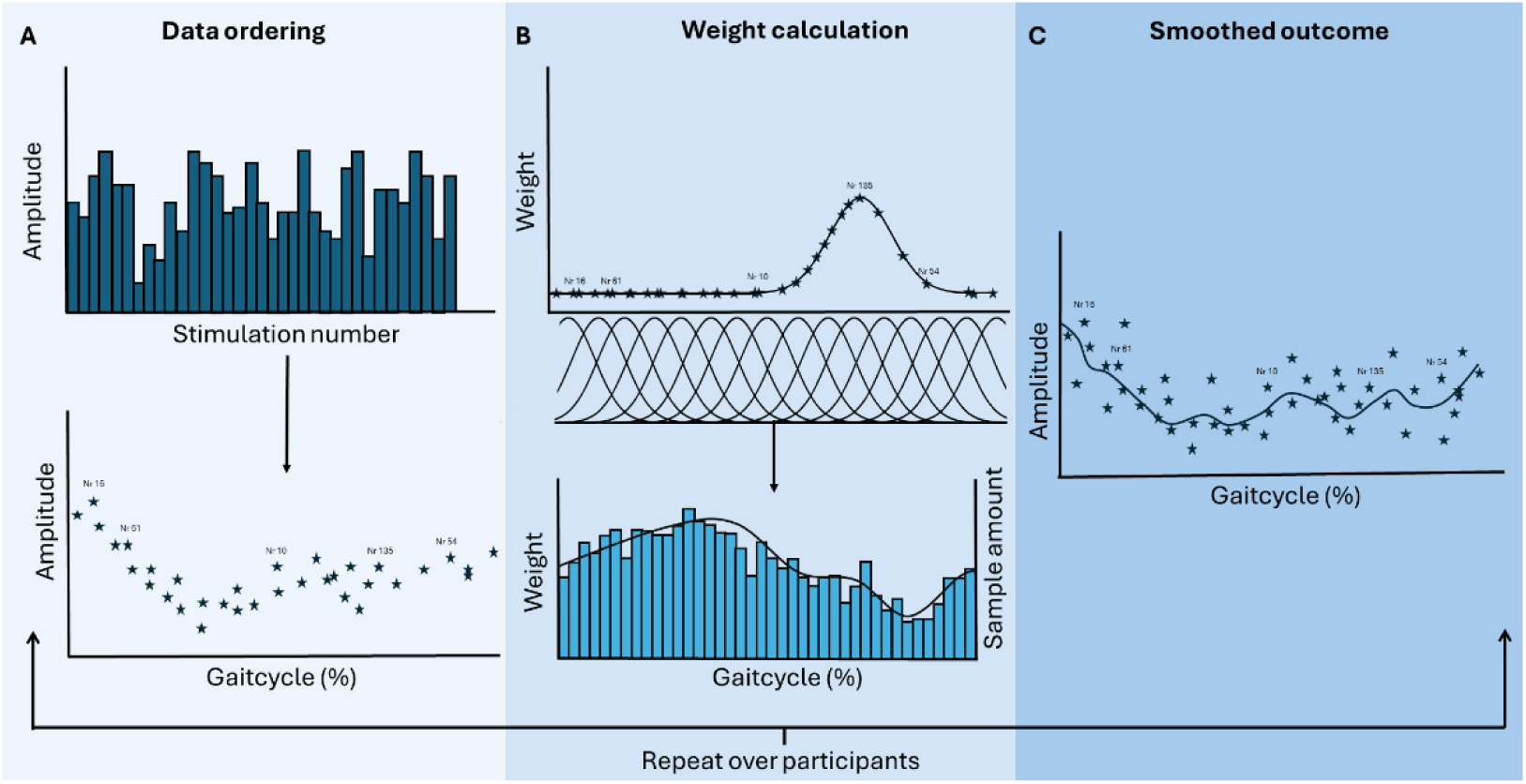
Illustration of the process of weighted smoothing for data from one condition of one participant. A) Stimulations and how the number of stimulation was linked to their gait cycle percentage. B) Display of how for each time sample the weight was distributed. Repeating this over the new temporal axis set at a chosen temporal resolution results in a general weight distribution based on sample density. C) The smoothed results from convolving the weight distribution with the stimulation amplitude.

To accommodate the cyclical nature of the data, the SMART code was modified by replacing the standard Gaussian distribution, which assumes linear data, with the Von Mises distribution (Supplementary data Figure 1). To prevent bias by outliers, a smoothing kernel containing sufficient data points was required. For this at least six data points had to be contained within 2 standard deviations (σ) of each weight distribution (Figure 3B and 4). As the data displayed an average density of 1.99 ± 0.58/1% (steady-state) and 2.02 ± 0.55/1% (laterally destabilised) for each participant, a σ of 2% was chosen. Consequently, the Von Mises distribution with a kappa (κ) of 60, which contained the same amount of datapoints as a normal distribution with a σ of 2% (Supplementary data Figure 1A), was used.

**Figure 4.**
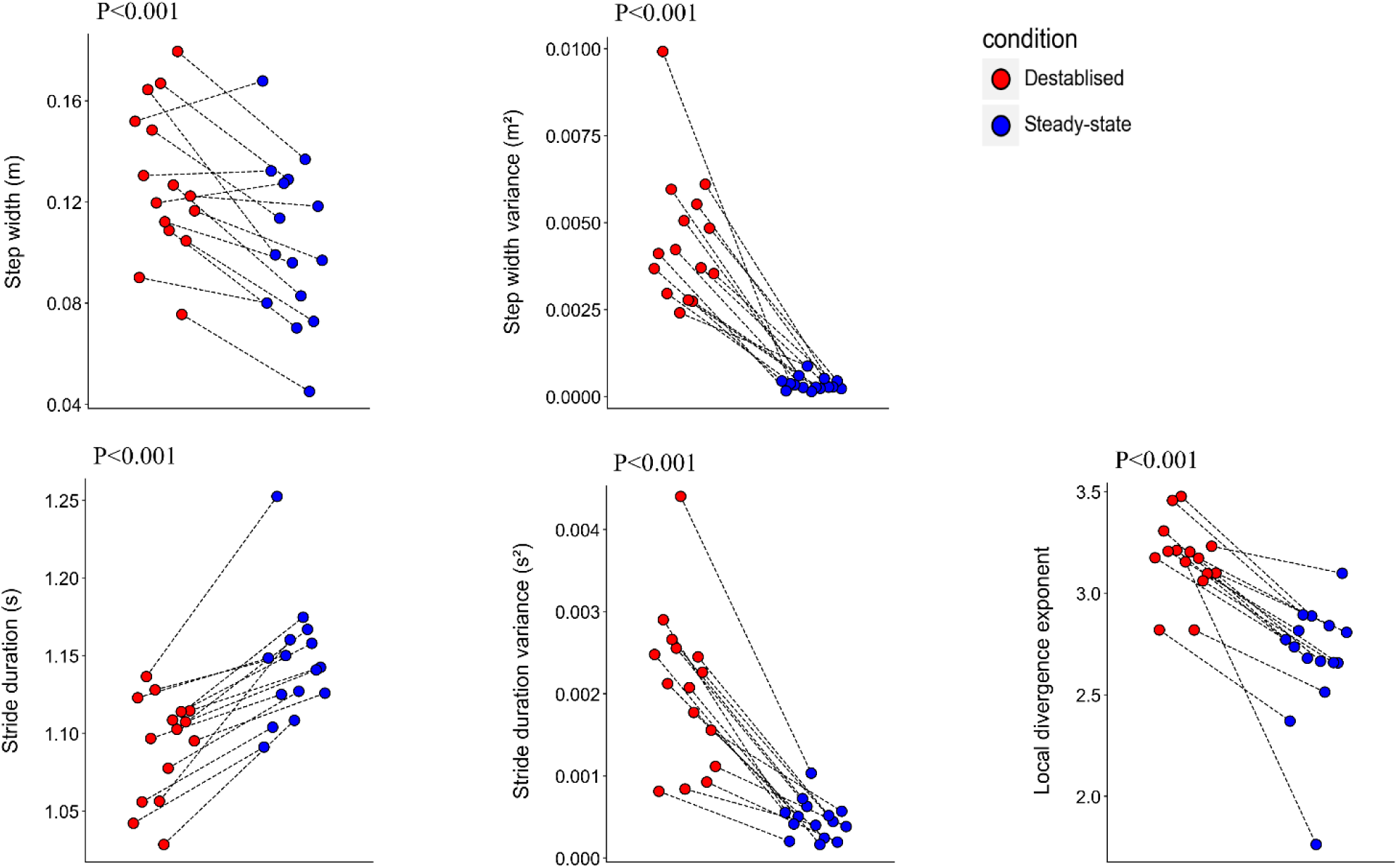
The effect of lateral destabilisation on step width, step width variance, stride duration, stride duration variance, and the local divergence exponent (λs). The dots represent the outcomes of individual participants, connected between conditions by black dotted lines to show the changes for each participant.

### Statistics

Statistics were performed in Python (version 3.12.0). All kinematic outcomes, apart from the relative explained variance of the foot placement model, were compared between steady-state and laterally destabilised gait using paired t-statistics.

The relative explained variance (R^2^) of the foot placement model over a step, ongoing EMG, absolute MEPs and MEP gains were compared between steady-state and laterally destabilised gait using paired cluster based-permutation testing according to the following procedure: First, paired-t-tests were applied to each time point, either through the weight-based approach, which computed the t-statistic based on the weighted average condition difference, or the non-weight based approach, utilising direct paired t-statistics. However, since each time point in a time series analysis is not independent of its neighbouring time points, condition differences were tested over clusters of consecutive time points, each of which exhibited statistically significant difference. Thus, cluster size was determined by the number of subsequent time points that are significantly different. This was done by summing the T values of a cluster and comparing this to summed T values of the permuted data’s cluster (i.e. shuffling the data in a random order followed by smoothing it). Clusters were considered significant if their summed T value was greater than the 95^th^ percentile of the 1000 permutations. This procedure was performed according to the code of Leeuwen and colleagues [60]. For all muscle activity and corticospinal excitability measures, this method was adjusted to account for the cyclic nature of the data, by wrapping the clusters around, connecting the clusters at the start and end of the gait cycle (https://doi.org/10.17605/OSF.IO/7AJUZ). For all statistical tests a significance threshold α of 0.05 was used.

## RESULTS

Of the 16 participants who completed the full experiment, one was excluded due to excessive relative movement between their head and the coil (i.e. fewer than 100 stimulations per condition remaining after exclusions). Data are presented from the remaining 15 participants (four males and 11 females; age 18–40 years). Seven participants performed the laterally destabilised condition first and seven performed the steady-state condition first. Two participants received TMS stimulation at 100% of motor threshold instead of the targeted 110% due to reporting discomfort at the higher intensity. However, MEPs were still observed in these participants and their results are included.

### Gait stability and quality of foot placement control

The lateral destabilisation evoked a significant increase in step width, step width variability, and stride duration variability and a significant decrease in stride duration and the local dynamic stability, demonstrated by increased λ_s_ (Figure 4).

The relative explained variance (Fisher-transformed R) of the foot placement model was greater in the laterally destabilised condition than in the steady-state condition at midstance. However, this difference was not observed in the rest of the stepcycle. Additionally, there was a significantly greater average foot placement error in the laterally destabilised condition (Figure 5).

**Figure 5.**
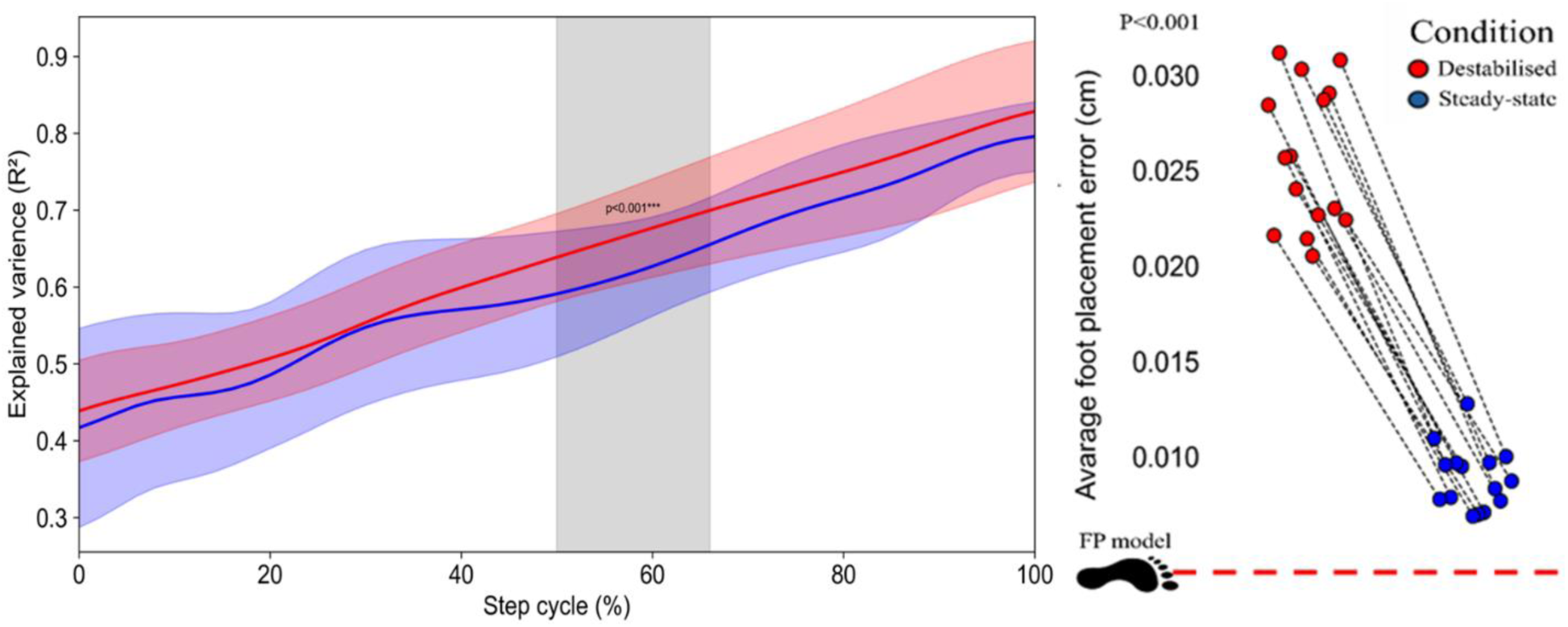
Explained variance of the foot placement as a function of step cycle. A step is defined from toe-off to subsequent heel strike. The coloured shaded area indicates the standard deviation of the corresponding condition. The grey shaded area indicate significant difference which was calculated based on the fishers transformed explained variance B) The average foot placement error (indicating how precise the actual foot placement is relative to the predicted foot placement) was increased during destabilised gait.

### Ongoing EMG and MEPs

In total, more than 3100 stimulations per condition (average 212 per participant in steady-state gait and 215 per participant in destabilised gait) were within the relative displacement threshold and included in analysis. This was 97.76% of the total stimulations (99.87% and 95.83% for steady-state and destabilised gait, respectively). For an overview of the coil displacement for each stimulation as a function of the gait cycle and condition, see Supplementary Figures 2 and 3. Steady-state and laterally destabilised gait were compared using a temporal resolution of 0.5% and a κ of 60. Our normalised weight distribution indicated standard deviations of sample spread of 7.7*10^-4^ and 4.9*10^-4^ for the steady-state and the laterally destabilised condition, respectively (Supplementary data Figure 4). The figures below display the results from the weight-based approach. All results from the non-weight-based approach can be found in the supplementary data (Supplementary data figure 5: ongoing EMG, 6: absolute MEPs, 7: MEP gains). Results were similar for both approaches, supporting the robustness of the findings.

#### Ongoing EMG

Muscle activity was greater during destabilised gait than during steady-state gait during some phase of the gait cycle in all muscles except the gastrocnemius medialis (Figure 6).

**Figure 6.**
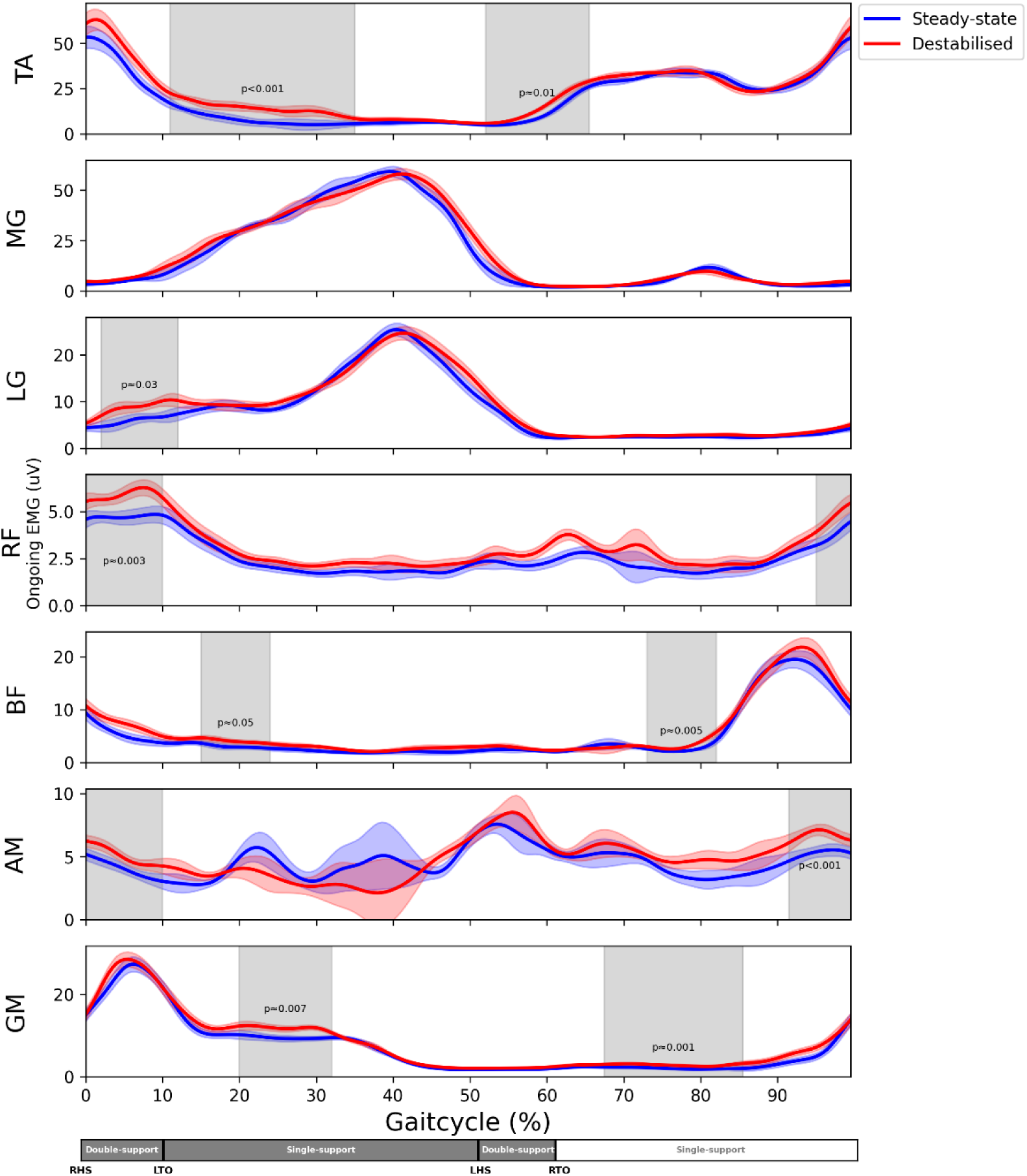
Ongoing EMG’s (uV) as function of the gait cycle for all muscles. 0% corresponds to right heel strike. The coloured shaded area’s are the standard deviations of the conditions. The grey shade around it indicate significant difference. M. gastrocnemius medialis (MG), m. gastrocnemius lateralis (LG), m. rectus femoris (RF), m. biceps femoris (BF), m. adductor magnus (AM), right heel strike (RHS), left heel strike (LHS), right toe off (RTO), left toe off (LTO).

#### Absolute MEPs

No differences in absolute MEP amplitude were detected in the lower-leg muscles (gastrocnemius medialis, gastrocnemius lateralis, and tibialis anterior). However, all upper-leg muscles demonstrated greater absolute MEPs during laterally destabilised gait than during steady-state gait (Figure 7). In the rectus femoris, MEP amplitude was larger over almost the entire gait cycle. A similar consistent increase in absolute MEPs was observed in the biceps femoris, with no differences during terminal swing and a short period of the stance phase. The adductor magnus exhibited increased absolute MEPs from terminal swing into the early stance phase. In the gluteus medius, three periods of significantly increased absolute MEPs were observed: during early stance, from late stance into early swing, and during the late swing phase.

**Figure 7.**
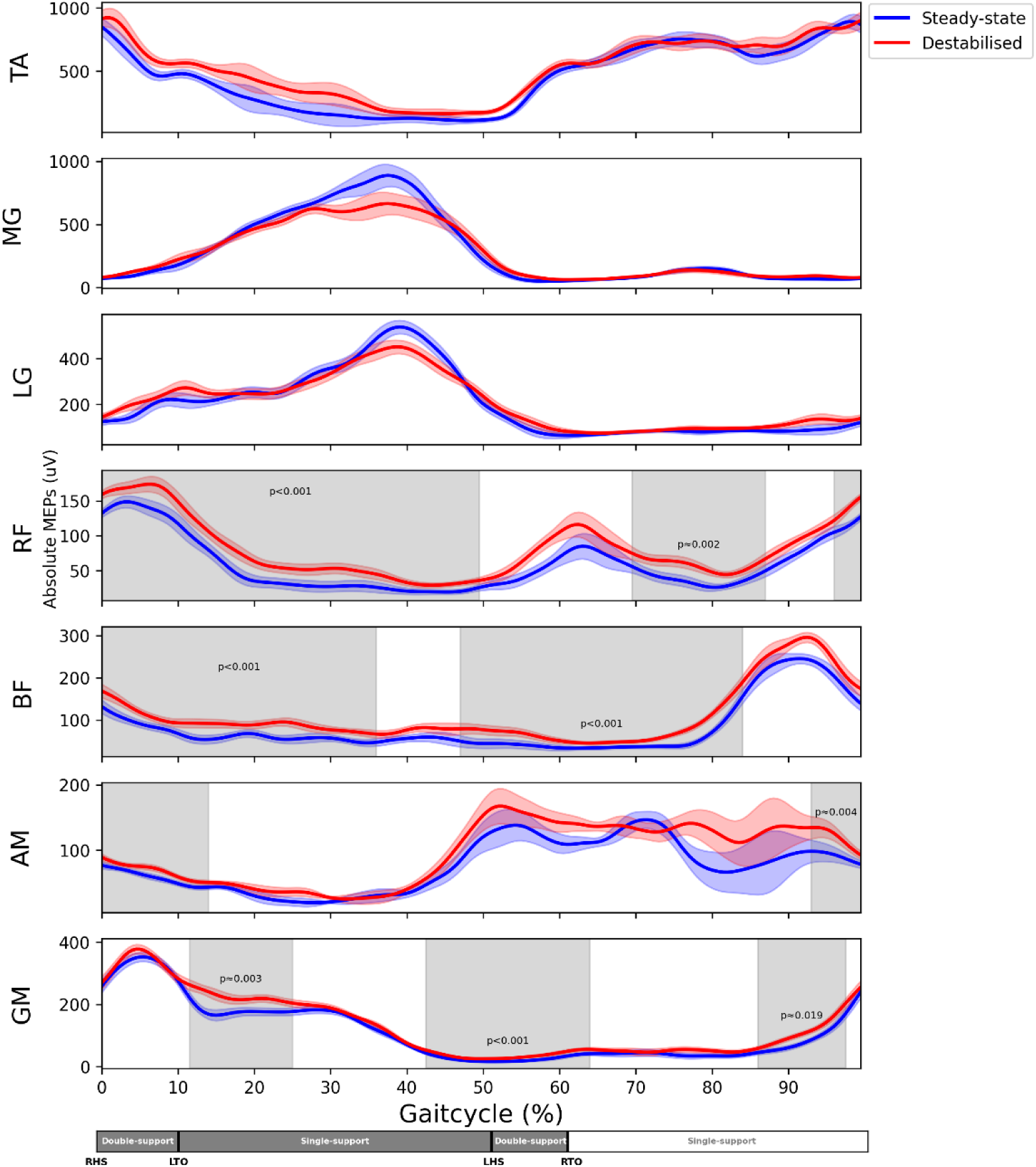
Absolute MEPs(uV) as function the gait cycle for all muscles. 0% corresponds to right heel strike. The coloured shaded area’s are the standard deviations of the conditions. The grey shade around it indicate significant difference. M. gastrocnemius medialis (MG), m. gastrocnemius lateralis (LG), m. rectus femoris (RF), m. biceps femoris (BF), m. adductor magnus (AM), right heel strike (RHS), left heel strike (LHS), right toe off (RTO), left toe off (LTO).

#### MEP gain

There was a short period of significantly increased MEP gain during destabilised gait in the tibialis anterior, and periods of increased MEP gain in all upper-leg muscles (Figure 8). In tibialis anterior, MEP gain was greater during laterally destabilised gait than during steady-state walking during the transition from single-leg stance to the double support phase. In the rectus femoris MEP gain was greater during laterally destabilised gait during the stance phase, and in the biceps femoris it was greater during the stance phase and from double support into mid-swing. The adductor magnus muscle exhibited two periods of significantly increased MEP gains, both during mid stance. Lastly, one period of increased MEP gain was observed in the gluteus medius during the second double support phase.

**Figure 8.**
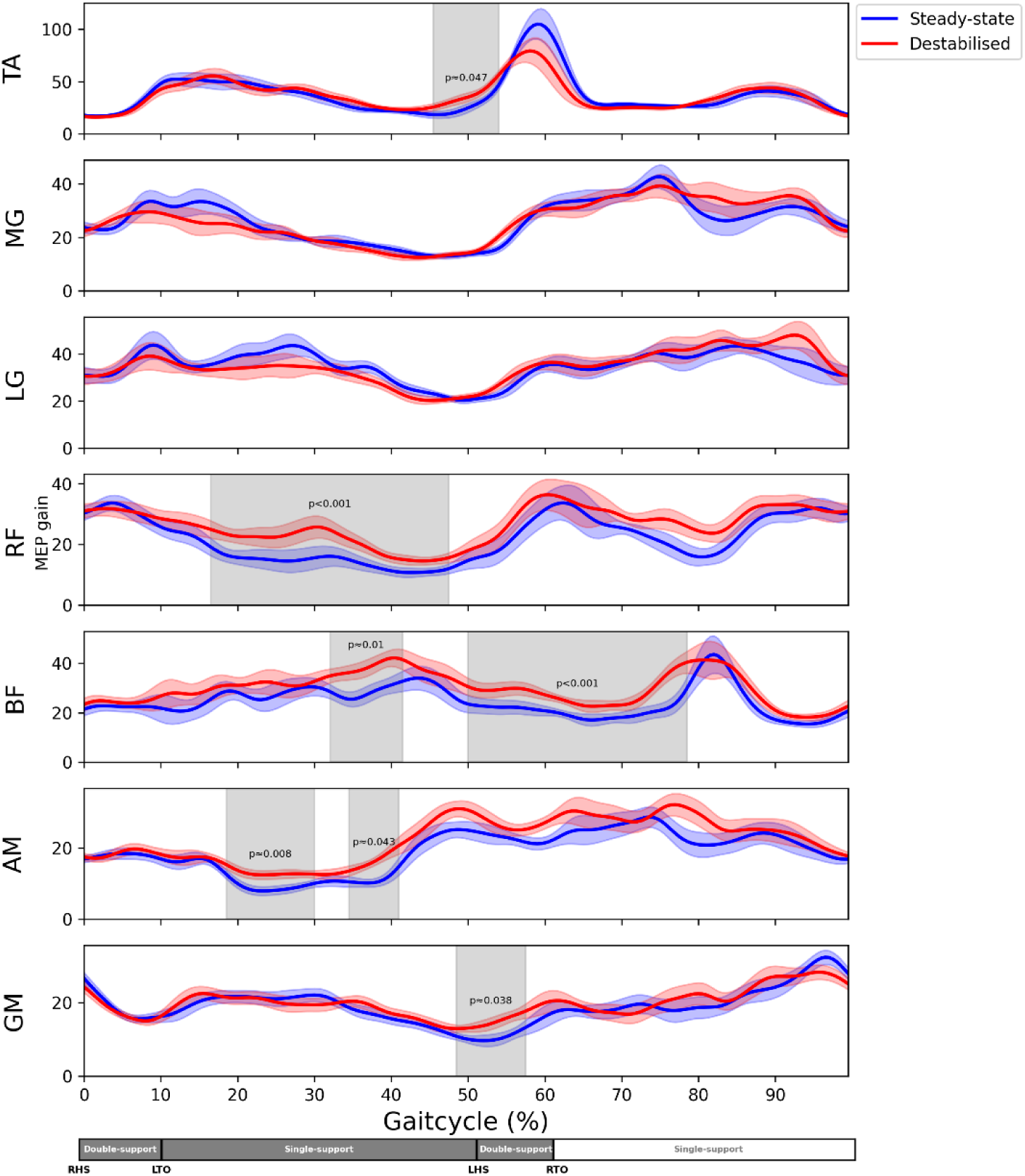
MEP gains are displayed over the gait cycle for all muscles. The plot starts at 0% of the gait cycle which corresponds to the right heel strike. The coloured shaded area’s are the standard deviations of the conditions. The grey shade around it indicate significant difference. M. gastrocnemius medialis (MG), m. gastrocnemius lateralis (LG), m. rectus femoris (RF), m. biceps femoris (BF), m. adductor magnus (AM), right heel strike (RHS), left heel strike (LHS), right toe off (RTO), left toe off (LTO).

## DISCUSSION

We investigated whether unpredictable mediolateral gait instability was associated with greater excitability of the corticospinal tract to multiple leg muscles. For this purpose, we compared MEPs elicited during steady-state treadmill gait to those elicited during laterally destabilised treadmill gait, where destabilisation was achieved using continuous pseudorandom mediolateral oscillations of the treadmill. This destabilisation increased gait instability and caused participants to adopt a more cautious walking strategy. This was accompanied by increased activity of proximal and distal leg muscles and increased excitability of the corticospinal tract, particularly to proximal leg muscles.

### Gait stability and quality of foot placement control

Similar to other studies [61,62], we observed a larger step width, step width variability, stride duration variability and shorter stride duration during laterally destabilised gait than during steady-state gait. Furthermore, we observed increased *λ_s_* in the laterally destabilised condition, indicating poorer local dynamic stability, i.e. more susceptibility to small perturbations [63]. Increased variability in stride duration and step width is often associated with increased risk of falls [43]; however, it can also be attributed to a purposeful adjustment response to overcome the stability demands [11,61]. One of the main mechanisms behind this adjustment is foot placement control. While we did observe a tightening of this foot placement control during midstance, this did not result in a tightening of the actual foot placement (i.e. reduction in foot placement error at the end of the step cycle). The quality of foot placement control can increase due to a greater variance in the CoM_states_, which most likely will happen due to the treadmill’s oscillations. Thus, we also estimated the foot placement control error [64], and found that this increased during lateral destabilised gait (Figure 4), indicating a decreased precision in foot placement. This likely results from the constant nature of the perturbation, as foot placement modulation requires time to execute [34,65,66], and from the type of perturbation, since medial surface translations are primarily managed through trunk-based strategies [67]. Altogether participants widened their steps and increased stride frequency to achieve the task in the presence of perturbations [29]. Increases in step width and reductions in step length are general strategies characterising “cautious walking” in unpredictable situations [29,68,69].

It is important to consider that the gait parameters measured during familiarisation might have deviated from those during the actual trial due to the introduction of TMS. TMS itself already works as a central perturbation, leading to anticipation and compensatory adjustments for the evoked movements [49]. While both conditions should be equally affected by the additional stimulation, the gait patterns during the familiarisation might not perfectly align those during the trials.

### Muscle activity and absolute MEPs

All stimulations were delivered at a location and intensity optimised for the tibialis anterior muscle. Although other muscles and stimulation sites could have been used to determine the motor threshold, the cortical representations of most leg muscles are located in close proximity, thus different localisation and intensity was not deemed necessary [52]. Muscle activity and absolute MEP amplitudes followed a similar phasic modulation over the gait cycle, which is in line with previous studies [17–19]. The cautious walking pattern employed in laterally destabilised gait was associated with greater activity of all muscles except gastrocnemius medialis. In the upper-leg muscles, this was mirrored by greater absolute MEPs. However, in lower-leg muscles there was no change in the amplitude of absolute MEPs. This was unexpected, but may be due to the method of destabilisation employed.

During steady-state gait, mediolateral stability is the dynamic equilibrium between the moments created by gravity and the forces generated by the muscles [70,71]. Primarily, the hip abductors, plantar flexors and to a lesser extent the dorsiflexors produce forces counteract the destabilisation due to gravity [23,70,72]. However, we suggest that in our laterally destabilised gait condition there was little demand on lower-leg muscles to compensate for the perturbations. Although lower-leg muscles play an important role in steady-state gait stability [73] and ankle moments have the capacity to correct for preceding foot placement errors [32], the unpredictability of constant and random surface shifts during gait may have necessitated a greater reliance on the upper-leg muscles. This is reflected by the rectus femoris and biceps femoris demonstrating increases in MEP amplitude over a large portion of the gait cycle, which likely served as a proactive control strategy [15], aligning with the ‘cautious walking’ strategy. For the gluteus medius and adductor magnus the responses seem more specific, with the increases aligning the muscles contributions to mediolateral acceleration [70]. Therefore, we suggest that the method of destabilisation limited the applicability of lower-leg muscles strategies, forcing participants to rely on upper-leg muscles to maintain gait stability.

### MEP gain

The clear link between ongoing (pre-stimulus) EMG and absolute MEP amplitude is consistent with the established relation between ongoing voluntary muscle activity and MEP amplitude [74,75]. This relation appears linear at low levels of activity (up to 10% of maximum) [74], but may be non-linear at higher contraction intensities [75]. This likely contributes to the lower MEP gains observed during more active phases of the gait cycle. However, this would also predict lower MEP gain in the destabilised walking, where muscle activity is greater, yet this was not observed.

Notably, a few muscles showed increases in MEP gain prior to the increase in muscle activity (Figure 8: tibialis anterior from 50 to 60%, biceps femoris 70 to 85% and gluteus medius 95 to 5%). These increases align with findings of Schubert and colleagues [21], who reported early increases in tibialis anterior and gastrocnemius medialis MEPs prior to phasic muscle activity changes, which they interpreted as a preparatory mechanism to preserve gait stability. In our study, the MEP gains in these periods were greater during laterally destabilised gait for the tibialis anterior and biceps femoris, although this effect of condition was not consistently observed across all muscles and did not occur in muscles known to be most involved in stabilising gait [11], such as the gluteus medius. Therefore, we suspect that an increase in this pre-activity MEP gain does not universally occur in response to externally imposed increases in stability demands. Further research is required to clarify the mechanisms behind the early facilitations.

Where there were difference in MEP gain across conditions, these were always in the direction of greater MEP gain during destabilised gait than during steady-state gait. No such differences were observed in the gastrocnemii, which likely reflects the limited use of ankle strategies to counteract our method of destabilisation [31,67,76]. The tibialis anterior showed greater MEP gain during the second double support phase in destabilised gait, which may serve to keep the forefoot elevated and prepare for impact [77,78]. The gluteus medius also showed only a brief increase in MEP gain near the second double support phase. Although this phase involves some transitional stabilisation demands and may also be linked to collision preparation and prevention [66,79], it is not the phase during which the gluteus medius has its greatest stabilisation potential, which occurs during mid-swing or mid-stance [11,34,66]. In general, the observed increases in MEP gain do not clearly align with specific stabilisation strategies [70,73], and there were no greater increases in muscles expected to play a dominant role in stability control, such as the gluteus medius. This raises the question of whether these differences in MEP gain between destabilised and steady-state gait reflect specific stability control mechanisms or rather general phasic postural adjustments.

We should take into account that our results could be affected by the inherent variability in both ongoing EMG signals and MEPs. EMG signals become more susceptible to noise at during periods of low muscle activity [80] and MEPs are subject to rapid, spontaneous fluctuations in corticospinal excitability, especially during periods of low excitability [81,82]. This may complicate the interpretation of MEP gains at low levels of muscle activity. However, the greater inter-stimulus variability in MEP amplitude at low activity levels may be expected to make it harder to detect differences between conditions. The observed differences in MEP gain were consistently in the direction of larger gain in destabilised gait, suggesting that this may not be driven by random variability. The non-linear relation between EMG and MEP amplitude, causing MEP gain to be lower at higher contraction intensities [74,75], may contribute to the absence of differences between conditions during highly active periods of the gait cycle.

Overall, we interpret these results as indicating increases in MEP gain during period of low muscle activity during laterally destabilised walking, particularly in proximal leg muscles, and suggest that this reflects changes in corticospinal excitability beyond simply reflecting ongoing muscle activity [50,74]. This may be related to increased task complexity or proactive stability control during destabilised gait. However, better methods to account for the influence of ongoing muscle activity on MEPs are needed to confirm this interpretation.

### Conclusions

In conclusion, we investigated the effects of unpredictable mediolateral instability on the excitability of the corticospinal tract to multiple leg muscles. We found that the imposed continuous unpredictable mediolateral movement of the treadmill increased instability and led participants to adopt a plethora of “cautious walking” strategies. This was associated with periods of increased absolute corticospinal excitability in all upper-leg muscles along with sustained increases in MEP gain in proximal upper-leg muscles, particularly at times of low muscle activity. These findings suggest that continuous unpredictable postural destabilisations heightened the responsiveness of the corticospinal tract to proximal leg muscles, suggesting that the corticospinal tract is involved in the adaptations to these conditions.

## ADDITIONAL INFORMATION

### Supplementary material

Opensource material of the raw data/codes and the questionnaire can be found at: https://doi.org/10.17605/OSF.IO/7AJUZ.

## REFERENCES

1. Kiehn, O. (2006). Locomotor circuits in the mammalian spinal cord. Annual Review of Neuroscience, 29, 279–306.

2. Fedirchuk, B., Nielsen, J., Petersen, N., & Hultborn, H. (1998). Pharmacologically evoked fictive motor patterns in the acutely spinalized marmoset monkey (*Callithrix jacchus*). Experimental Brain Research, 122, 351–361.

3. Liddell, E. G. T., & Phillips, C. G. (1944). Pyramidal section in the cat. Brain, 67, 1–9.

4. Shik, M. L., & Orlovsky, G. N. (1976). Neurophysiology of locomotor automatism. Physiological Reviews, 56, 465–501.

5. Duysens, J., & Van de Crommert, H. W. A. A. (1998). Neural control of locomotion; Part 1: The central pattern generator from cats to humans. Gait & Posture, 7, 131–141.

6. Muir, G. D., & Whishaw, I. Q. (1999). Complete locomotor recovery following corticospinal tract lesions: Measurement of ground reaction forces during overground locomotion in rats. Behavioural Brain Research, 103, 45–53.

7. Dietz, V. (1992). Human neuronal control of automatic functional movements: Interaction between central programs and afferent input. Physiological Reviews, 72, 33–69.

8. Pribe, C., Grossberg, S., & Cohen, M. A. (1997). Neural control of interlimb oscillations. Biological Cybernetics, 77, 141–152.

9. Knutsson, E., & Richards, C. (1979). Different types of disturbed motor control in gait of hemiparetic patients. Brain, 102, 405–430.

10. Barthélemy, D., Grey, M. J., Nielsen, J. B., & Bouyer, L. (2011). Involvement of the corticospinal tract in the control of human gait (pp. 181–197).

11. Bruijn, S. M., & van Dieën, J. H. (2018). Control of human gait stability through foot placement. Journal of the Royal Society Interface, 15, 20170816.

12. Siebner, H. R., Funke, K., Aberra, A. S., Antal, A., Bestmann, S., Chen, R., et al. (2022). Transcranial magnetic stimulation of the brain: What is stimulated? – A consensus and critical position paper. Clinical Neurophysiology, 140, 59–97.

13. Di Lazzaro, V., & Rothwell, J. C. (2014). Corticospinal activity evoked and modulated by non-invasive stimulation of the intact human motor cortex. Journal of Physiology, 592, 4115– 4128.

14. Groppa, S., Schlaak, B. H., Münchau, A., Werner-Petroll, N., Dünnweber, J., Bäumer, T., et al. (2012). The human dorsal premotor cortex facilitates the excitability of ipsilateral primary motor cortex via a short latency cortico-cortical route. Human Brain Mapping, 33, 419–430.

15. Bestmann, S., & Krakauer, J. W. (2015). The uses and interpretations of the motor-evoked potential for understanding behaviour. Experimental Brain Research, 233, 679–689.

16. Hupfeld, K. E., Swanson, C. W., Fling, B. W., & Seidler, R. D. (2020). TMS-induced silent periods: A review of methods and call for consistency. Journal of Neuroscience Methods, 346, 108950.

17. Capaday, C., Lavoie, B. A., Barbeau, H., Schneider, C., & Bonnard, M. (1999). Studies on the corticospinal control of human walking. I. Responses to focal transcranial magnetic stimulation of the motor cortex. Journal of Neurophysiology, 81, 129–139.

18. Petersen, T. H., Willerslev-Olsen, M., Conway, B. A., & Nielsen, J. B. (2012). The motor cortex drives the muscles during walking in human subjects. Journal of Physiology, 590, 2443–2452.

19. Barthélemy, D., & Nielsen, J. B. (2010). Corticospinal contribution to arm muscle activity during human walking. Journal of Physiology, 588, 967–979.

20. Mate, K. K. V. (2022). Transcranial magnetic stimulation during gait. Neurology India, 70, 1448–1453.

21. Schubert, M., Curt, A., Jensen, L., & Dietz, V. (1997). Corticospinal input in human gait: Modulation of magnetically evoked motor responses. Experimental Brain Research, 115, 234–246.

22. Roelker, S. A., Kautz, S. A., & Neptune, R. R. (2019). Muscle contributions to mediolateral and anteroposterior foot placement during walking. Journal of Biomechanics, 95, 109310.

23. Neptune, R. R., & McGowan, C. P. (2016). Muscle contributions to frontal plane angular momentum during walking. Journal of Biomechanics, 49, 2975–2981.

24. Hof, A. L. (2007). The equations of motion for a standing human reveal three mechanisms for balance. Journal of Biomechanics, 40, 451–457.

25. Bauby, C. E., & Kuo, A. D. (2000). Active control of lateral balance in human walking. Journal of Biomechanics, 33, 1433–1440.

26. O’Connor, S. M., & Kuo, A. D. (2009). Direction-dependent control of balance during walking and standing. Journal of Neurophysiology, 102, 1411–1419.

27. Hurt, C. P., Rosenblatt, N., Crenshaw, J. R., & Grabiner, M. D. (2010). Variation in trunk kinematics influences variation in step width during treadmill walking by older and younger adults. Gait & Posture, 31, 461–464.

28. Wang, Y., & Srinivasan, M. (2014). Stepping in the direction of the fall: The next foot placement can be predicted from current upper body state in steady-state walking. Biology Letters, 10, 20140405.

29. Perry, J. A., & Srinivasan, M. (2017). Walking with wider steps changes foot placement control, increases kinematic variability and does not improve linear stability. Royal Society Open Science, 4, 160627.

30. Arvin, M., van Dieën, J. H., & Bruijn, S. M. (2016). Effects of constrained trunk movement on frontal plane gait kinematics. Journal of Biomechanics, 49, 3085–3089.

31. Hof, A. L., & Duysens, J. (2018). Responses of human ankle muscles to mediolateral balance perturbations during walking. Human Movement Science, 57, 69–82.

32. van Leeuwen, A. M., van Dieën, J. H., Daffertshofer, A., & Bruijn, S. M. (2021). Ankle muscles drive mediolateral center of pressure control to ensure stable steady state gait. Scientific Reports, 11, 21481.

33. Rankin, B. L., Buffo, S. K., & Dean, J. C. (2014). A neuromechanical strategy for mediolateral foot placement in walking humans. Journal of Neurophysiology, 112, 374–383.

34. Hof, A. L., & Duysens, J. (2013). Responses of human hip abductor muscles to lateral balance perturbations during walking. Experimental Brain Research, 230, 301–310.

35. van Leeuwen, A. M., van Dieën, J. H., Daffertshofer, A., & Bruijn, S. M. (2020). Active foot placement control ensures stable gait: Effect of constraints on foot placement and ankle moments. PLoS ONE, 15, e0242215.

36. Schrager, M. A., Kelly, V. E., Price, R., Ferrucci, L., & Shumway-Cook, A. (2008). The effects of age on medio-lateral stability during normal and narrow base walking. Gait & Posture, 28, 466–471.

37. Osoba, M. Y., Rao, A. K., Agrawal, S. K., & Lalwani, A. K. (2019). Balance and gait in the elderly: A contemporary review. Laryngoscope Investigative Otolaryngology, 4, 143–153.

38. Skiadopoulos, A., Moore, E. E., Sayles, H. R., Schmid, K. K., & Stergiou, N. (2020). Step width variability as a discriminator of age-related gait changes. Journal of NeuroEngineering and Rehabilitation, 17, 41.

39. Callisaya, M. L., Blizzard, L., Schmidt, M. D., McGinley, J. L., & Srikanth, V. K. (2010). Ageing and gait variability--a population-based study of older people. Age and Ageing, 39, 191–197.

40. Schulz, B., Bruns, M., Mierau, A., & Strüder, H. K. (2012). Changes in intracortical inhibition of the primary motor cortex in elderly adults and their relationship to manual dexterity. Age, 34, 1313–1329.

41. Maki, B. E., & McIlroy, W. E. (1996). Postural control in the older adult. Clinics in Geriatric Medicine, 12, 635–658.

42. Brauer, S. G., Burns, Y. R., & Galley, P. (2000). A prospective study of laboratory and clinical measures of postural stability to predict community-dwelling fallers. Journal of Gerontology A: Biological Sciences and Medical Sciences, 55(12), M469–M476.

43. Toebes, M. J. P., Hoozemans, M. J. M., Furrer, R., Dekker, J., & van Dieën, J. H. (2012). Local dynamic stability and variability of gait are associated with fall history in elderly subjects. Gait & Posture, 36(4), 527–531.

44. Brach, J. S., Berlin, J. E., VanSwearingen, J. M., Newman, A. B., & Studenski, S. A. (2005). Too much or too little step width variability is associated with a fall history in older persons who walk at or near normal gait speed. Journal of NeuroEngineering and Rehabilitation, 2, 21.

45. Hausdorff, J. M., Rios, D. A., & Edelberg, H. K. (2001). Gait variability and fall risk in community-living older adults: A 1-year prospective study. Archives of Physical Medicine and Rehabilitation, 82(8), 1050–1056.

46. Schubert, M., Curt, A., Colombo, G., Berger, W., & Dietz, V. (1999). Voluntary control of human gait: Conditioning of magnetically evoked motor responses in a precision stepping task. Experimental Brain Research, 126(4), 583–588.

47. Dambreville, C., Neige, C., Mercier, C., Blanchette, A. K., & Bouyer, L. J. (2022). Corticospinal excitability quantification during a visually-guided precision walking task in humans: Potential for neurorehabilitation. Neurorehabilitation and Neural Repair, 36(9), 689–700.

48. Bonnard, M., Camus, M., Coyle, T., & Pailhous, J. (2002). Task-induced modulation of motor evoked potentials in upper-leg muscles during human gait: A TMS study. European Journal of Neuroscience, 16(12), 2225–2230.

49. Camus, M., Pailhous, J., & Bonnard, M. (2006). On-line flexibility of the cognitive tuning of corticospinal excitability: A TMS study in human gait. Brain Research, 1076(1), 144–149.

50. Nandi, T., Fisher, B. E., Hortobágyi, T., & Salem, G. J. (2018). Increasing mediolateral standing sway is associated with increasing corticospinal excitability, and decreasing M1 inhibition and facilitation. Gait & Posture, 60, 135–140.

51. Rossi, S., Hallett, M., Rossini, P. M., & Pascual-Leone, A. (2011). Screening questionnaire before TMS: An update. Clinical Neurophysiology, 122(8), 1686.

52. Davies, J. L. (2020). Using transcranial magnetic stimulation to map the cortical representation of lower-limb muscles. Clinical Neurophysiology Practice, 5, 87–99.

53. Dingwell, J. B., & Cusumano, J. P. (2000). Nonlinear time series analysis of normal and pathological human walking. Chaos, 10(4), 848–863.

54. Dingwell, J. B., & Marin, L. C. (2006). Kinematic variability and local dynamic stability of upper body motions when walking at different speeds. Journal of Biomechanics, 39(3), 444–452.

55. Rosenstein, M. T., Collins, J. J., & De Luca, C. J. (1993). A practical method for calculating largest Lyapunov exponents from small data sets. Physica D: Nonlinear Phenomena, 65(1-2), 117–134.

56. Stenum, J., Bruijn, S. M., & Jensen, B. R. (2014). The effect of walking speed on local dynamic stability is sensitive to calculation methods. Journal of Biomechanics, 47(16), 3776– 3779.

57. Bruijn, S. M. (2017). Local dynamic stability. Zenodo.

58. Afschrift, M., Bruijn, S., & van Dieen, J. H. (2023). Assessment of stabilizing feedback control of walking, a tutorial.

59. Yang, F., & Pai, Y.-C. (2014). Can sacral marker approximate center of mass during gait and slip-fall recovery among community-dwelling older adults? Journal of Biomechanics, 47(17), 3807–3812.

60. van Leeuwen, J., Smeets, J. B. J., & Belopolsky, A. V. (2019). Forget binning and get SMART: Getting more out of the time-course of response data. *Attention, Perception*, & Psychophysics, 81(10), 2956–2967.

61. McAndrew, P. M., Dingwell, J. B., & Wilken, J. M. (2010). Walking variability during continuous pseudo-random oscillations of the support surface and visual field. Journal of Biomechanics, 43(8), 1470–1475.

62. Onushko, T., Boerger, T., Van Dehy, J., & Schmit, B. D. (2019). Dynamic stability and stepping strategies of young healthy adults walking on an oscillating treadmill. PLoS One, 14(3), e0212207.

63. Bruijn, S. M., Bregman, D. J. J., Meijer, O. G., Beek, P. J., & van Dieën, J. H. (2012). Maximum Lyapunov exponents as predictors of global gait stability: A modelling approach. Medical Engineering & Physics, 34(4), 428–436.

64. Mahaki, M., van Leeuwen, A. M., Bruijn, S. M., van der Velde, N., & van Dieën, J. H. (2023). Mediolateral foot placement control can be trained: Older adults learn to walk more stable, when ankle moments are constrained. PLoS One, 18(3), e0292449.

65. Vlutters, M., Van Asseldonk, E. H. F., & van der Kooij, H. (2018). Foot placement modulation diminishes for perturbations near foot contact. Frontiers in Bioengineering and Biotechnology, 6, 89.

66. Afschrift, M., Pitto, L., Aerts, W., van Deursen, R., Jonkers, I., & De Groote, F. (2018). Modulation of gluteus medius activity reflects the potential of the muscle to meet the mechanical demands during perturbed walking. Scientific Reports, 8(1), 11675.

67. Brough, L. G., & Neptune, R. R. (2024). A comparison of the effects of mediolateral surface and foot placement perturbations on balance control and response strategies during walking. Gait & Posture, 108, 313–319.

68. Wu, M., Matsubara, J. H., & Gordon, K. E. (2015). General and specific strategies used to facilitate locomotor maneuvers. PLoS One, 10(6), e0132707.

69. Hak, L., Houdijk, H., Steenbrink, F., Mert, A., van der Wurff, P., Beek, P. J., & van Dieën, J. H. (2013). Stepping strategies for regulating gait adaptability and stability. Journal of Biomechanics, 46(5), 905–911.

70. Pandy, M. G., Lin, Y.-C., & Kim, H. J. (2010). Muscle coordination of mediolateral balance in normal walking. Journal of Biomechanics, 43(11), 2055–2064.

71. MacKinnon, C. D., & Winter, D. A. (1993). Control of whole body balance in the frontal plane during human walking. Journal of Biomechanics, 26(6), 633–644.

72. Neptune, R. R., & McGowan, C. P. (2011). Muscle contributions to whole-body sagittal plane angular momentum during walking. Journal of Biomechanics, 44(1), 6–12.

73. Anderson, F. C., & Pandy, M. G. (2003). Individual muscle contributions to support in normal walking. Gait & Posture, 17(2), 159–169.

74. Darling, W. G., Wolf, S. L., & Butler, A. J. (2006). Variability of motor potentials evoked by transcranial magnetic stimulation depends on muscle activation. Experimental Brain Research, 174(3), 376–385.

75. Devanne, H., Lavoie, B. A., & Capaday, C. (1997). Input-output properties and gain changes in the human corticospinal pathway. Experimental Brain Research, 114(2), 329–338.

76. Koch, M., Eckardt, N., Zech, A., & Hamacher, D. (2020). Compensation of stochastic time-continuous perturbations during walking in healthy young adults: An analysis of the structure of gait variability. Gait & Posture, 80, 253–259.

77. Perera, C. K., Gopalai, A. A., Ahmad, S. A., & Gouwanda, D. (2021). Muscles affecting minimum toe clearance. Frontiers in Public Health, 9, 645576.

78. von Tscharner, V., Goepfert, B., & Nigg, B. M. (2003). Changes in EMG signals for the muscle tibialis anterior while running barefoot or with shoes resolved by non-linearly scaled wavelets. Journal of Biomechanics, 36(8), 1169–1176.

79. Ciunelis, K., Borkowski, R., & Błażkiewicz, M. (2024). The impact of induced acceleration perturbations in selected phases of the gait cycle on kinematic and kinetic parameters. Applied Sciences, 14(9), 4849.

80. De Luca, C. J. (1997). The use of surface electromyography in biomechanics. Journal of Applied Biomechanics, 13(2), 135–163.

81. iers, L., Cros, D., Chiappa, K. H., & Fang, J. (1993). Variability of motor potentials evoked by transcranial magnetic stimulation. Electroencephalography and Clinical Neurophysiology, 89(6), 415–423.

82. Cuypers, K., Thijs, H., & Meesen, R. L. J. (2014). Optimization of the transcranial magnetic stimulation protocol by defining a reliable estimate for corticospinal excitability. PLoS One, 9(7), e86380.

