## Supplementary figures for "Changes in Corticospinal Excitability in Response to Mediolateral Gait Instability"

### Supplementary data

Figure 1

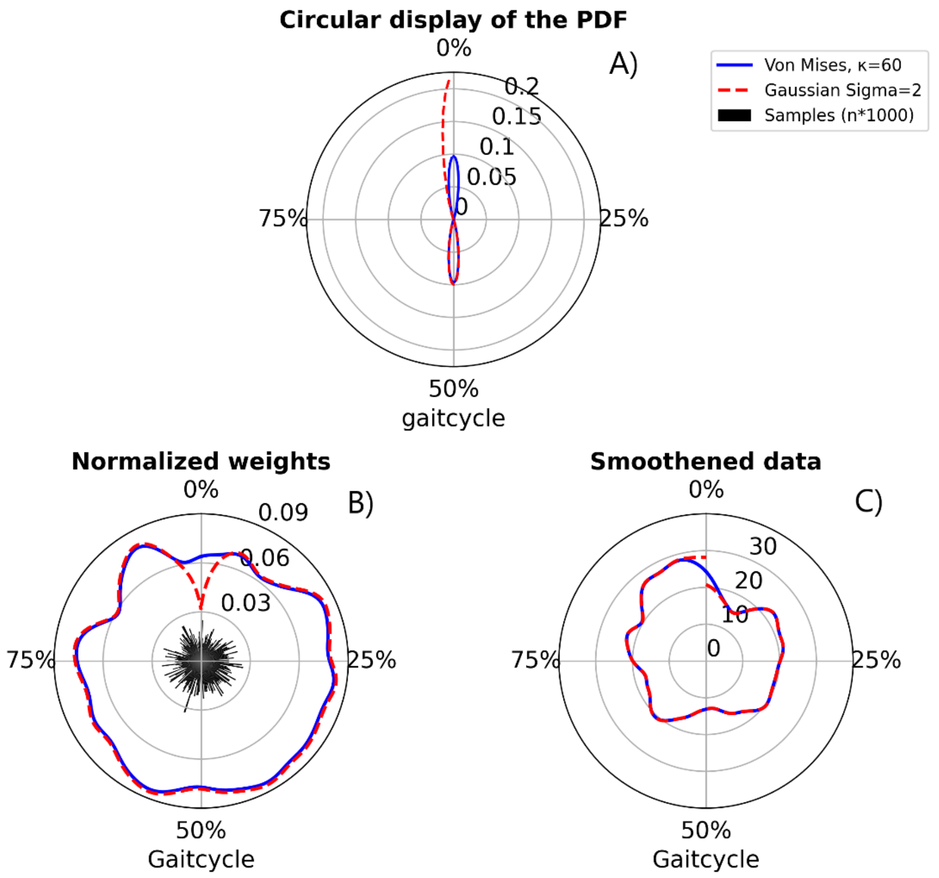


***Figure 1.*** *Smoothing differences between the Von Mises and Gaussian distributions.* ***A)*** *Differences at timepoints 0 and 50.0% for the probability density function (PDF) of both the Von Mises and Gaussian distribution.* ***B)*** *Effect of sample density on temporal weights and how this differs between linear and circular distributions. The sample density is displayed in black bars with the amount of samples per 0.5%. The amount if equal to the y-axis times 1000* ***C)*** *Effect of the difference in weight distribution on the smoothed data outcome.*

Figure 2

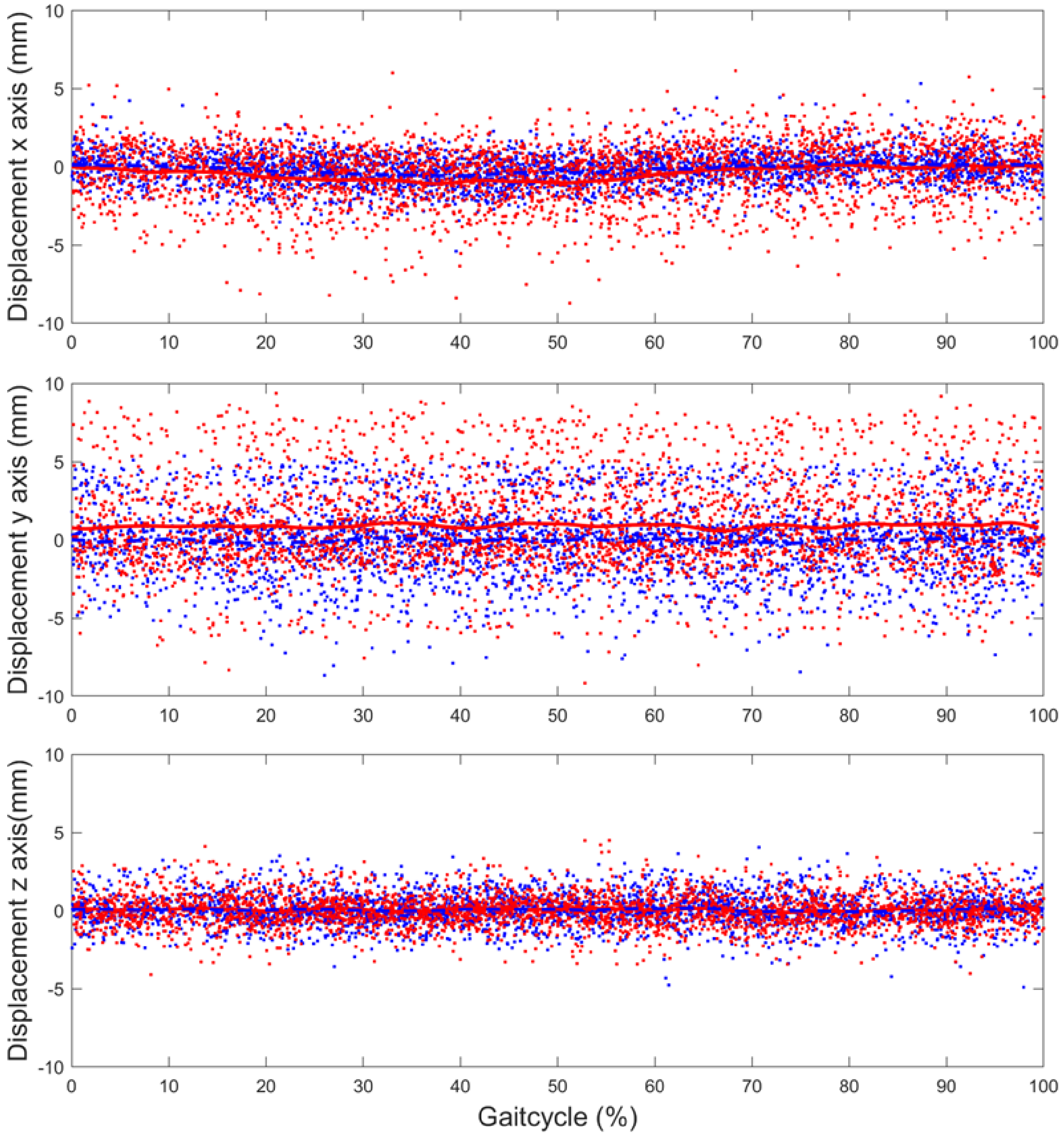


***Figure 2.*** *Display of displacement over the gait cycle. To indicate any phase dependent effects.*

Figure 3

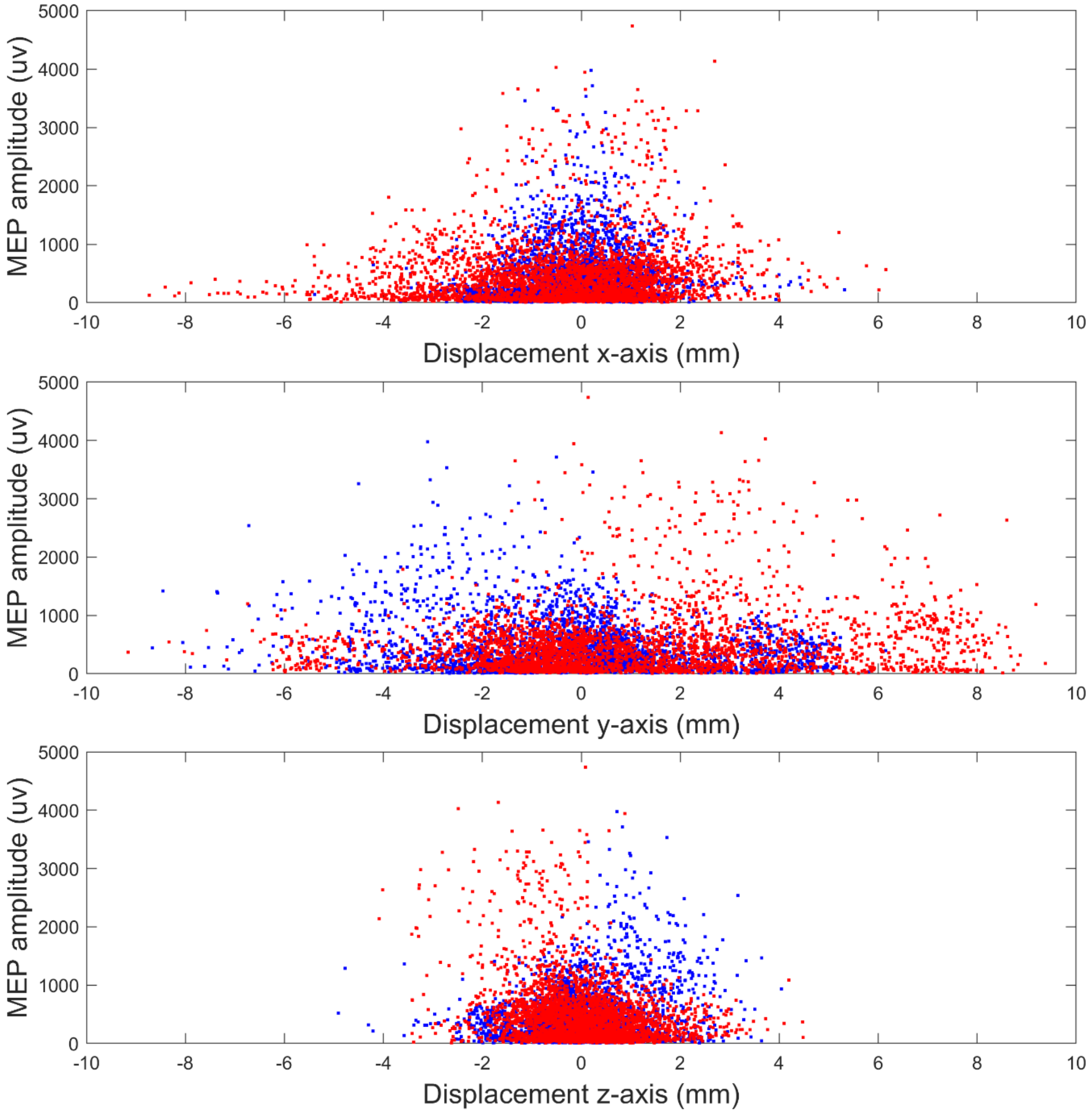


***Figure 3.*** *Display of the effects of displacement on MEP amplitudes for the tibialis anterior.*

Figure 4

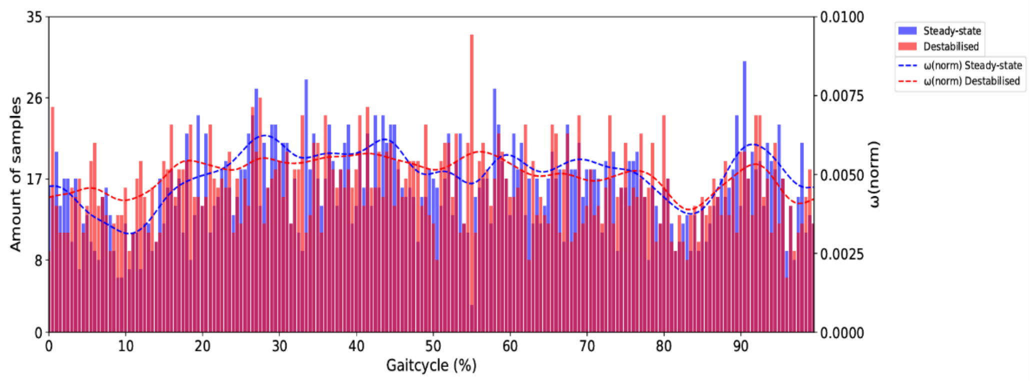


***Figure 4.*** *The amount samples at each temporal point (i.e. the amount of samples per 0.5%) summed over all participants. Purple is seen due to the overlapping bars of both conditions. The dotted lines indicate the normalised weight at each timepoint. This was calculated based on the sample density, a perfect homogeneous data spread would result in a flat line at 0.0050.*

Figure 5

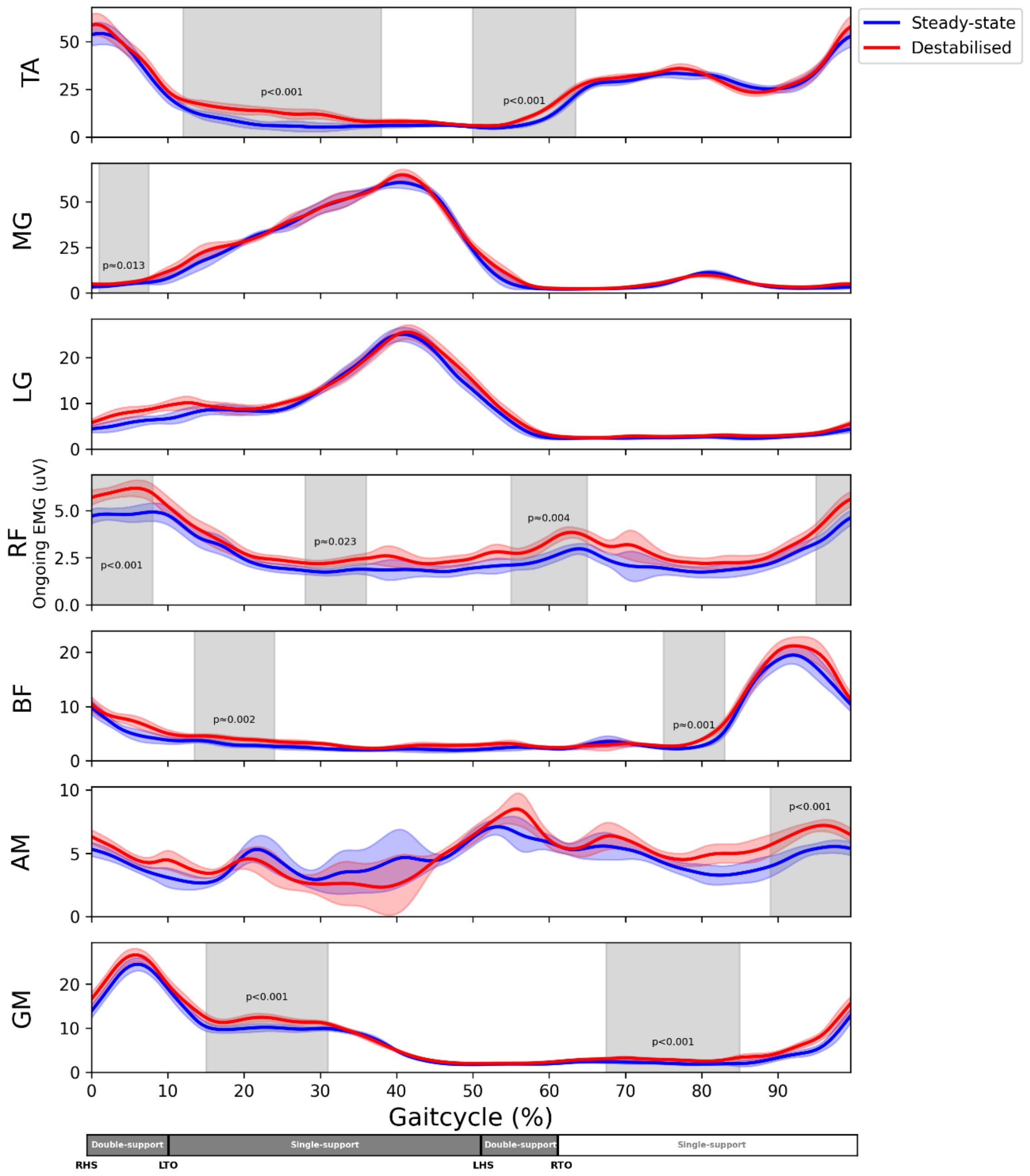


*__Figure 5.__ Non-weight-based statistics ongoing EMG's (uV) as function of the gait cycle for all muscles. 0% corresponds to right heel strike. The coloured shaded area's are the standard deviations of the conditions. The grey shade around it indicate significant difference. M. gastrocnemius medialis (MG), m. gastrocnemius lateralis (LG), m. rectus femoris (RF), m. biceps femoris (BF), m. adductor magnus (AM), right heel strike (RHS), left heel strike (LHS), right toe off (RTO), left toe off (LTO).*

Figure 6

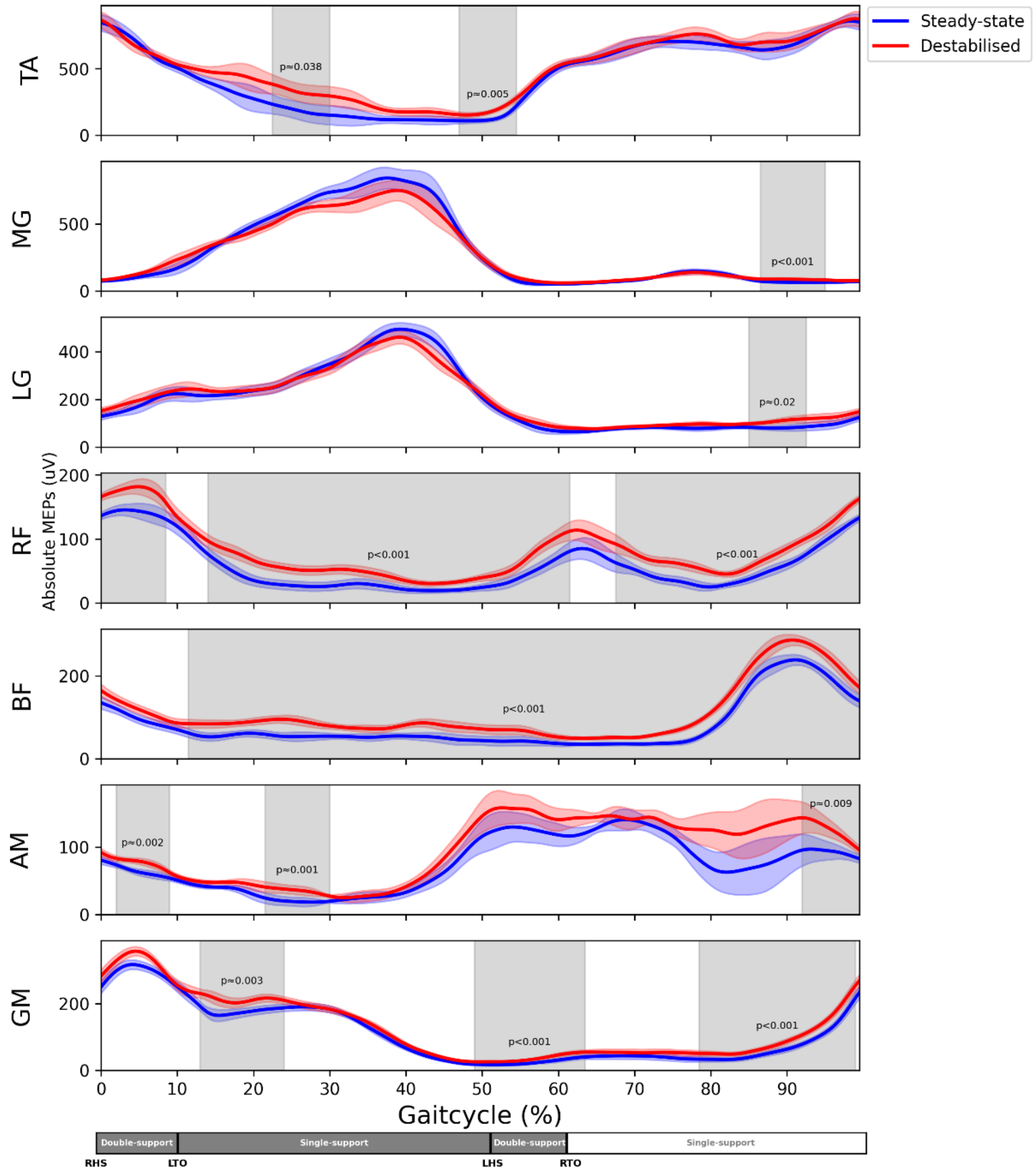


***Figure 6.*** *Non-weight-based statistics absolute MEPs(uV) as function the gait cycle for all muscles. 0% corresponds to right heel strike. The coloured shaded area's are the standard deviations of the conditions. The grey shade around it indicate significant difference. M. gastrocnemius medialis (MG), m. gastrocnemius lateralis (LG), m. rectus femoris (RF), m. biceps femoris (BF), m. adductor magnus (AM) , right heel strike (RHS), left heel strike (LHS), right toe off (RTO), left toe off (LTO).*

Figure 7

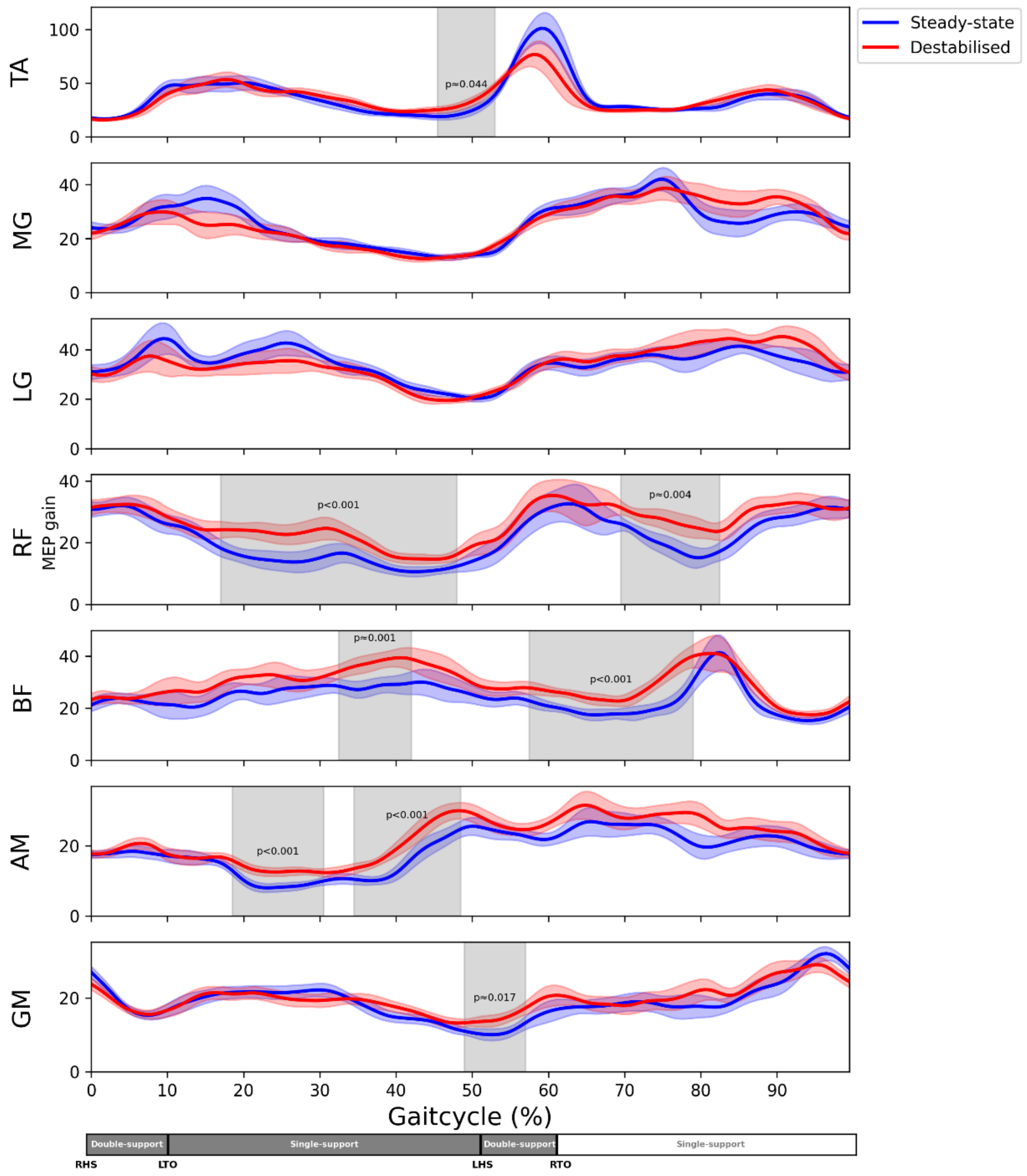


***Figure 7.*** *Non-weight-based statistics* MEP gains *are displayed over the gait cycle for all muscles. The plot starts at 0% of the gait cycle which corresponds to the right heel strike. The coloured shaded area's are the standard deviations of the conditions. The grey shade around it indicate significant difference. M. gastrocnemius medialis (MG), m. gastrocnemius lateralis (LG), m. rectus femoris (RF), m. biceps femoris (BF), m. adductor magnus (AM) , right heel strike (RHS), left heel strike (LHS), right toe off (RTO), left toe off (LTO).*
